# Variability in Individual Native Fibrin Fiber Mechanics

**DOI:** 10.1101/2024.07.29.605650

**Authors:** Christine C. Helms

## Abstract

Fibrin fibers are important structural elements in blood coagulation. They form a mesh network that acts as a scaffold and imparts mechanical strength to the clot. A review of published work measuring the mechanics of fibrin fibers reveals a range of values for fiber extensibility. This study investigates fibrinogen concentration as a possible variable responsible for variability in fibrin fiber mechanics. It expands previous work to describe the modulus, strain hardening, extensibility, and the force required for fiber failure when fibers are formed with different fibrinogen concentrations. Lateral force atomic force microscopy was used to create stress-strain curves for individual nanofibers and data was obtained from fibers formed from 0.5 NIH U/ml thrombin, 55 Loewy U/ml FXIII, and 1 mg/ml or 2 mg/ml fibrinogen. Analysis of the mechanical properties showed fiber formed from 1 mg/ml fibrinogen and 2 mg/ml fibrinogen had significantly different mechanical properties. To help clarify our findings we developed two behavior profiles to describe individual fiber mechanics. The first describes a fiber with low initial modulus and high extensible, that undergoes strain hardening around 100 % strain, and has moderate strength. A majority of fibers formed with 1 mg/ml fibrinogen showed this behavior profile. The second profile describes a fiber with a high initial modulus, minimal strain hardening, high strength, and low extensibility. Most fibrin fibers formed with 2 mg/ml fibrinogen were described with this second behavior profile. In conclusion, we see a range of behaviors from fibers formed from native fibrinogen molecules but various fibrinogen concentrations. Potential differences in fiber formation is investigated with SEM. It is likely this range of behaviors also occurs in vivo. Understanding the variability in mechanical properties could contribute to a deeper understanding of pathophysiology of coagulative disorders.

## 1. Introduction

Fibrinogen is an important plasma protein that plays a critical role in blood coagulation. During coagulation, fibrinogen is cleaved by thrombin to produce fibrin. Fibrin can self-assemble into two molecules-wide protofibrils known for their half-staggered arrangement. These protofibrils elongate and laterally aggregate into larger fibrin fibers which maintain the repetitive structure of the protofibrils. Electron microscopy, x-ray scattering, and atomic force microscopy have provided evidence of this organized structure within individual fibrin fibers [1-5]. Fibrin fibers branch to form a 3-D fiber network that is remarkably porous and plays a structural and mechanical role in blood clots. In addition to individual fibers, fiber can bundle together to form thick fibers [6-8] and more recently, the ability of fibrin to form fibrin sheets or film at the air-liquid interface has been reported and investigated [9,10]. These fibrin sheets may play a role in preventing bacterial invasion [11,12]. These sheets can tear and gather to form fibrin bundles resembling fibrin fibers [9].

There are still unanswered questions regarding the formation of fibrin assemblies, including an extensive description of lateral aggregation of protofibrils, mechanisms driving fiber bundling, and the molecular structure in fibrin films. These details of molecular interactions and fiber packing are directly related to the mechanical properties of the resultant structure. Therefore, measuring fiber mechanics can provide insight into molecular interactions and highlight structural differences. We aim to measure individual fiber mechanical properties to gain insight into the variability or lack thereof in molecular interactions in fibrin fibers and fibrin constructs. Furthermore, we seek to understand the mechanical properties of fibrin constructs since changes in fibrin properties directly relate to health and disease.

The mechanical properties of fibrin clots are contingent upon their network structure and the mechanics of their individual fibers. Alterations affecting the mechanical properties of fibrin clots can arise from changes at various scales, including individual molecules [13-16], individual fibers [17-19], interactions between fibers [6,20,21], interactions between fibers and their surroundings [22-24], and the overall network structure [25-27]. Measurements of clot mechanics have been found to correlate with reduced lysis, strength, and disease, highlighting the clinical significance of understanding fibrin mechanics [28-30].

Diseases related to thrombosis and hemostasis, such as cardiovascular disease, stroke, and myocardial infarction are correlated with fibrin mechanics. For example, Collet et al. showed increased clot stiffness and altered network structure in young post-myocardial infarction patients [31] and Martinez et al showed an accelerated growth of viscoelastic properties in pulmonary embolism compared to deep vein thrombosis patients [32]. Additionally, clot mechanics have been useful in disease assessment in sepsis [33].

Many studies have looked at environmental influences on fibrin clot structure and mechanics. Elements such as salt, calcium, protein post-translational modification and buffers have all been shown to effect clot properties [10,34,35]. Additionally, connections between protein concentrations and whole clot properties have been explored [36-38]. However, it is difficult to separate the effects of network structure from changes in individual fiber properties when studying the whole clot. Isolating the role of each component of clot mechanics provides greater insight and information about the mechanisms responsible for the macroscopic mechanics and may provide insight into fiber formation.

Measurements of individual fiber mechanics have grown during the past two decades. At the individual fiber level, fiber diameter has been identified as playing a crucial role in fiber mechanics [17]. It has been observed that as fiber diameter decreases, the modulus of individual fibrin fibers increases, attributed to dense molecular packing at the core of fibrin fibers [18,39]. Other studies have compared the mechanics of fibers formed from various fibrinogen molecules, such as native fibrinogen, fibrinogen variants, mutated recombinant fibrinogen molecules, and fibrinogen from different species [13,15,40]. These studies have highlighted the role of the aC-domain in extensibility, toughness, and modulus [15,41]. Additionally, they have indicated a role for gamma’ and gamma:gamma crosslinking in fiber packing and mechanics [16].

Beyond the effect of fiber diameter, few studies have investigated variability in individual native fibrin fiber mechanics. To our knowledge, each previous study of individual fibrin mechanics used a single set of protein concentration. However, comparison between experiments reveals variations in reported fiber mechanics, specifically fiber extensibilities in purified systems showed trends when plotted verse fibrinogen concentration or thrombin to fibrinogen ratio (Fig. 1).

**Fig 1.**
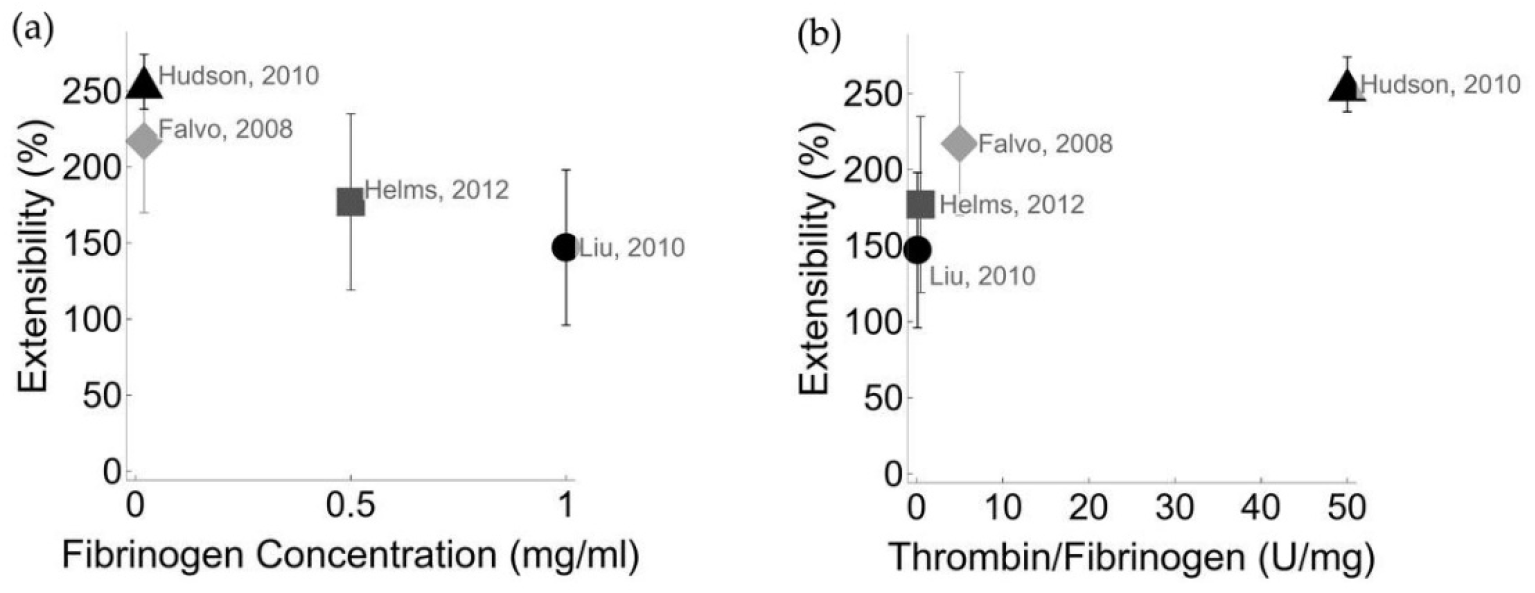
The extensibility of fibrin fibers reported in the literature [13,15,21,42]. (a) Extensibility data is plotted versus fibrinogen concentration. (b) Extensibility is plotted versus thrombin to fibrinogen ratio. All data are from fibers formed from purified fibrinogen and crosslinked with factor XIIIa.

In this work we investigate variations in individual fiber mechanics due to fibrinogen concentration. We identify a range of fiber mechanical behaviors and develop behavior profiles that describe the properties of two distinct sets of fiber behavior.

## 2. Methods

### 2.1 Substrate preparation

Patterned substrates for fibrin atomic force manipulation were prepared as previously described [41,42]. Briefly, a micropatterned PDMS stamp was created using SYLGARD 184 Silicone Elastomer and an etched silicon master wafer. This stamp was pressed into UV-curing glue (Norland 81 optical adhesive, Jamesburg, NJ, USA) on top of a #1.5 cover glass slide then cured and removed leaving behind a patterned substrate of repeating 10 micrometer wide ridges and 12 μm wide and 7 μm deep wells.

### 2.2 Fibrin Fiber formation

Fibrin clots were formed in a buffer of 10 mM HEPES (pH 7.4), 5 mM CaCl_2_ and 140 mM NaCl, which we will refer to as fibrin buffer. Clots were formed with a final concentration of 1 mg/ml or 2 mg/ml fibrinogen, unless otherwise indicate, 55 Lowey U/ml FXIII and 0.1 U/ml thrombin. All proteins were acquired from Enzyme Research laboratories (South Bend, Indiana, USA). Fibrin, FXIII, and buffer were mixed in a microtube. Thrombin was added and the solutions was immediately mixed via pipetting and moved to the patterned substrate. The solution was given between 2 and 4 hours to coagulate in a humid environment. After coagulation excess clot was removed by gently pulling a pipette tip over the surface of the clot and followed by rinsing the substrate 3 times with calcium-free fibrin buffer. 20 nm carboxylate-modified fluorescent microspheres (FluoSpheres, Invitrogen, Waltham, MA, USA) were diluted to a concentration of 10^5^ per ml in calcium-free fibrin buffer. 100 μl of the diluted microspheres was placed on top of the sample for 10 minutes before the sample was rinse with buffer to remove excess beads. The samples were then stored in a pool of buffer until manipulation by the AFM. All AFM manipulations were performed within 10 hours of sample formation.

### 2.3 Atomic Force Microscope Lateral Force Manipulation

Individual fiber mechanics were determined using a combined atomic force microscope (AFM) and fluorescence microscope procedure as previously described [15,42-44]. An AFM cantilever (CSC38 Tip B, NanoandMore, Watsonville, CA) was calibrated for lateral force measurements. The lateral force is directly related to the torque of the cantilever.

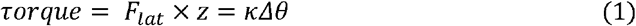

Because Δ*θ* is small, we can approximate the arc length traveled by the point of the tip with the linear displacement Δ*x*.

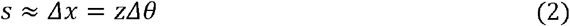

The distance z, from the point of rotation to the application of the force depends on the height of the tip, h, the thickness of the cantilever, t, and the distance from the point of the tip to the location of force, d (often 2 μm, Fig.2).

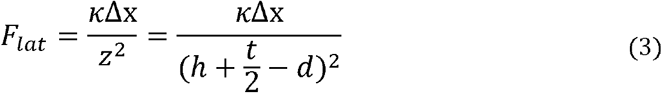

**Fig. 2.**
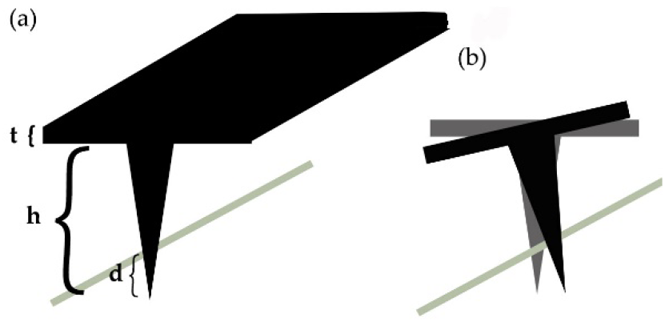
A schematic of the cantilever pressing into a fiber (green). (a) The height, cantilever thickness and distance form the point of the tip to the location of the force are indicated. (b) The force on the cantilever due to the fiber causes the tip to rotate from its original position shown in gray to its rotated position show in black.

Next using Euler-Lagrange beam mechanics and correcting for length of the cantilever to the tip, the torsion constant can be found by

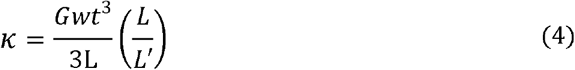

where G is the shear modulus of the cantilever, w is the width, L is the length, and L’ is the length to the tip. Normal force resonance was used to determine the thickness. Substituting into the equation for lateral force allows us to define a *k*_*lat*_.

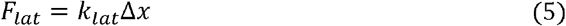

Where *k*_*lat*_ is the lateral spring constant of the cantilever and Δ*x* is the displacement of the cantilever tip. The relationship between the tip displacement and lateral voltage signal provided by the AFM software was measured by pushing the tip into a glass edge and obtaining a lateral voltage versus tip displacement curve.

The movement of the cantilever tip was controlled with a script written in the lab for an Asylum MFP-3D-BIO AFM. The script allows the operator of the AFM to select a step size, direction, and stepping rate for the cantilever. The fibers were manipulated at a strain rate of 150 nm/s.

### 2.4 Scanning Electron Microscopy

After manipulation samples were prepared for scanning electron microscopy (SEM, JEOL 6360 LV SEM, Peabody, MA, USA). First samples were fixed with 2.5 % glutaraldehyde mixed with fibrin buffer and allowed to crosslink for 30 minutes. Then the samples were thoroughly rinsed and placed in a series of ethanol dilutions starting at 30 % ethanol and ending at 100 % ethanol. Next the samples were critical point dried, and sputter coated using a gold-palladium target. Lastly, samples were mounted and imaged at 10 kV and magnifications between 20,000x and 23,000x.

### 2.5 Statistical Analysis

Data are presented as the mean and standard deviation of the mean. Comparisons were preformed using Student’s t-test and ANOVA. Data were considered statistically significant when the p < 0.5. Excel and Mathematica software were used for statistical analysis.

## 3. Results

### 3.1 Fibrin to Thrombin ratio

The ratio of thrombin to fibrinogen is important in clot characterization as it helps govern clotting kinetics and therefore, has been reported in previous publications [2,38,45]. We analyzed the extensibility data from the literature with respect to the ratio of thrombin to fibrinogen as well as fibrinogen concentration. We found that an increase in fibrinogen concentration led to a decrease in extensibility and a decrease thrombin to fibrin ratio led to a decrease in extensibility, as seen in Fig. 1.

To replicate the data from the literature we took measurements of fibrin extensibility while adjusting the thrombin to fibrinogen ratio. A range of fibrinogen and thrombin concentrations were used with thrombin ranging from 0.05 U/ml to 1 U/ml and fibrinogen ranging from 0.5 mg/ml to 2 mg/ml. When we aggregated the data using the thrombin to fibrinogen ratio, we could not discern a clear trend (Fig. 3a). This is likely because the effects of changing thrombin and changing fibrinogen are not proportional [38]. However, when we considered the data using the concentrations of thrombin and fibrinogen a trend became apparent. Fibers formed with low protein concentrations, for example 0.1 U/ml thrombin and 0.1 mg/ml fibrinogen, had a lower average extensibility than fibers formed from higher protein concentrations, 1 U/ml thrombin and 1 mg/ml fibrinogen. Fig. 3b shows this trend for the ratios of 1 U/mg and 2 U/mg thrombin to fibrinogen. For example, at a ratio of 2 U/ml thrombin to fibrinogen when the concentration of thrombin and fibrinogen are both less than 0.1 the extensibility is very low.

**Fig. 3.**
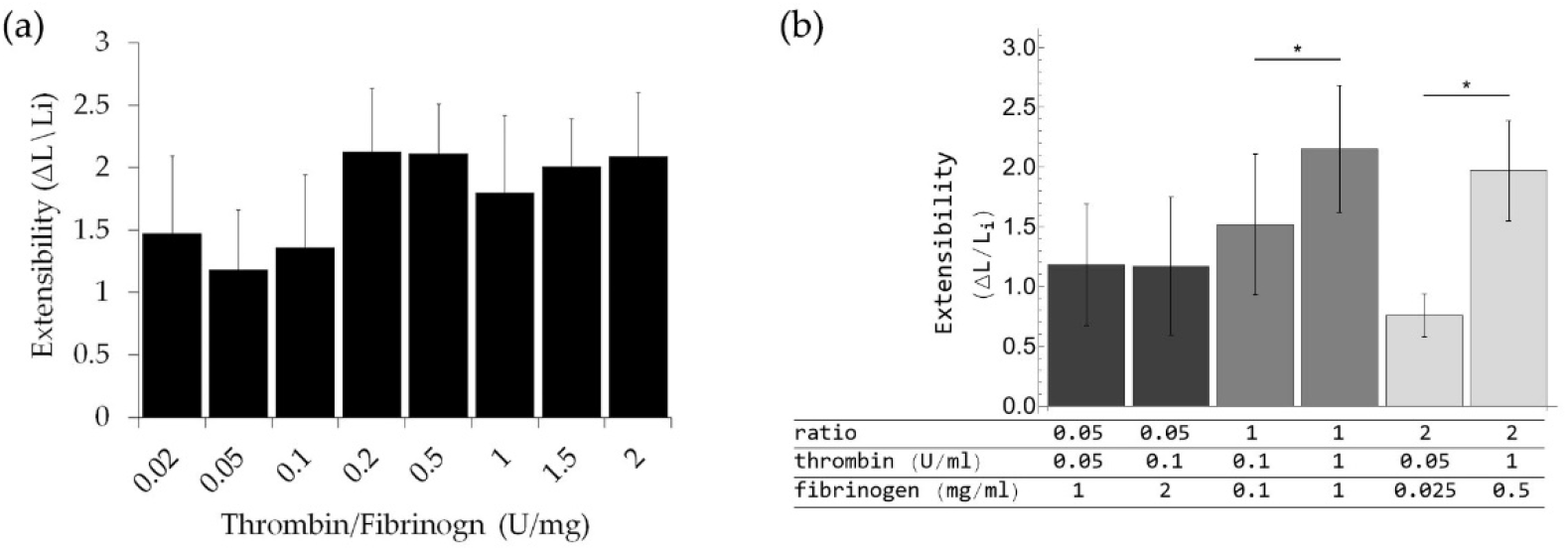
(a) A plot of the average fiber extensibility versus thrombin to fibrinogen ratio. Each bar represernts and aggregate of various concentrations of fibrinogen and thrombin producing the same protein ratio. (b) A plot of the average fiber extensibility versus thrombin to fibrin ratio. In this plot when various thrombin or fibrin concentrations are used the data is kept separate. At a ratio of 1 and 2, we see significant increase in extensibility when the protein concentrations are higher (* p < 0.01)

To simplify the system, we choose to hold thrombin concentration constant and vary fibrinogen concentration. Fibers were formed from a range of fibrinogen concentrations, 0.1 mg/ml, 0.5 mg/ml, 1 mg/ml, 1.5 mg/ml, and 2 mg/ml. Fig. 4 shows the extensibility had a maximum at the intermediate fibrinogen concentration of 1 mg/ml. Fibers formed at the lowest and highest concentrations tested, 0.1 mg/ml and 2 mg/ml, had significantly lower extensibilities. The largest difference in extensibility was seen between the 1 mg/ml fibrinogen fibers and the 2 mg/ml fibrinogen fibers. Therefore, these concentrations were chosen for further investigation of individual fiber mechanics.

**Fig. 4b.**
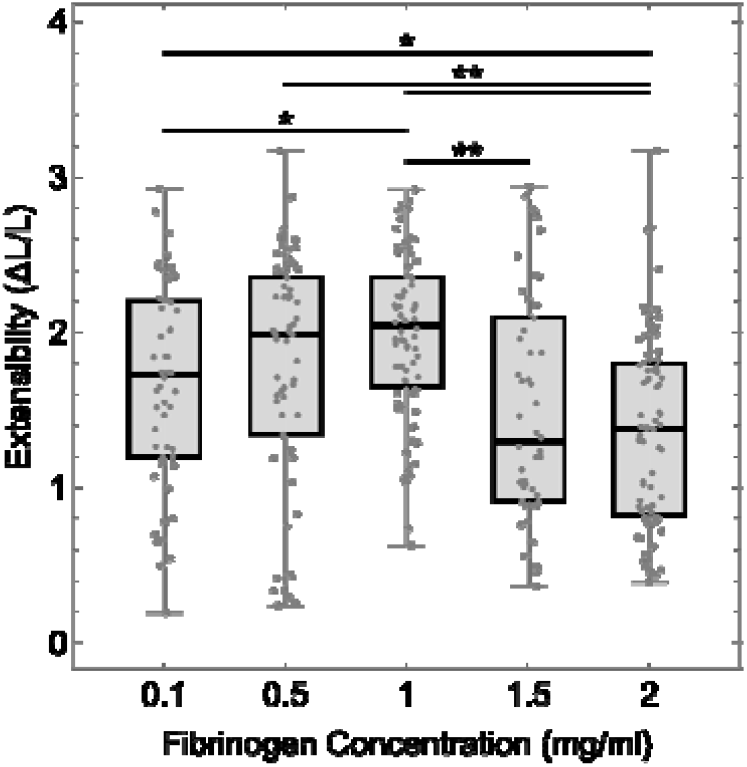
A box and whisker plot displaying fibrin fiber extensibility versus fibrinogen concentration. At each concentration at least 50 fibers were tested. The box displays the 75 % quantile, the 25 % quantile and the median, while the whiskers display the upper and lower fence (** p-value < 0.001, * p-value < 0.01).

### 3.2 Fibers formed with 1 mg/ml fibrinogen and 2 mg/ml fibrinogen

We obtained force verse strain curves for 33 fibers. Analysis of the force verse strain curves suggested two distinct profiles of fiber behavior. One behavior profile described fibers that were initially soft, underwent significant strain hardening and had large extensibilities. The other behavior profile described fibers that were initially stiff, only slightly increased their modulus during manipulation, and had lower extensibility. The observed mechanical behavior related to fibrinogen concentration during fiber formation. Our data and analysis focus on fibers formed from 1 mg/ml fibrinogen and 2 mg/ml fibrinogen.

#### 3.2.1 Fiber diameter by SEM

Fiber diameters were obtained by SEM. At the completion of AFM testing the samples were submerged in a series of ethanol dilutions, critical point dried, and sputter coated. Fiber diameters of fibers formed from 1 mg/ml and 2 mg/ml fibrinogen were 112 ± 5 nm (n = 32, mean ± standard deviation of the mean) and 101 ± 4 nm (n = 41), respectively and were not significantly different (p > 0.05). This agrees with work by Belcher et al. where fiber diameter increased between 1 mg/ml and 5 mg/ml fibrinogen at low thrombin concentrations. However the increase in diameter was within one standard deviation [46]. The average diameters obtained by SEM were used to convert the AFM force data into modulus data. Fibers are comprised of approximately 70 % water and 30 % protein therefore drying steps needed for SEM diameter measurements may cause the measured fiber diameters to differ from hydrated fibrin diameters. However, work by Belcher et al. showed SEM fiber diameters and hydrated fibrin diameters measured by super resolution fluorescence microscopy agree within 10%, suggesting minimal effects from drying [46]. Either way, uncertainty in fiber diameter is important to consider when interpreting the reported modulus values. An underestimate of the average fiber diameter by a given percent will lead to an overestimate of their modulus by double that percent. While the values for modulus may contain systematic uncertainty due to diameter measurements, comparisons between the two fiber populations should be unaffected.

#### 3.2.2 Fiber appearance by SEM

There were differences in appearance of fibers formed from the different fibrin concentrations. Samples formed from 2 mg/ml contained fibers that spanned the trough and continued onto the ridge, similar to Fig. 5a, as well as fibers with “root” structures attaching them to the ridge or a “film-like” structure at the ridge (Fig. 6). This root and film like structure has been attributed to fibers formed from fibrin film [9]. Fibers with root or films structures were often seen in densely populated areas of the sample. These densely populated areas are avoided during AFM manipulation because lateral manipulation requires adequate space between fibers. Samples formed from 1 mg/ml fibrinogen were dominated by fibers that continued onto the ridge and only contained a few fibers that showed dense branching near ridge.

**Fig. 5.**
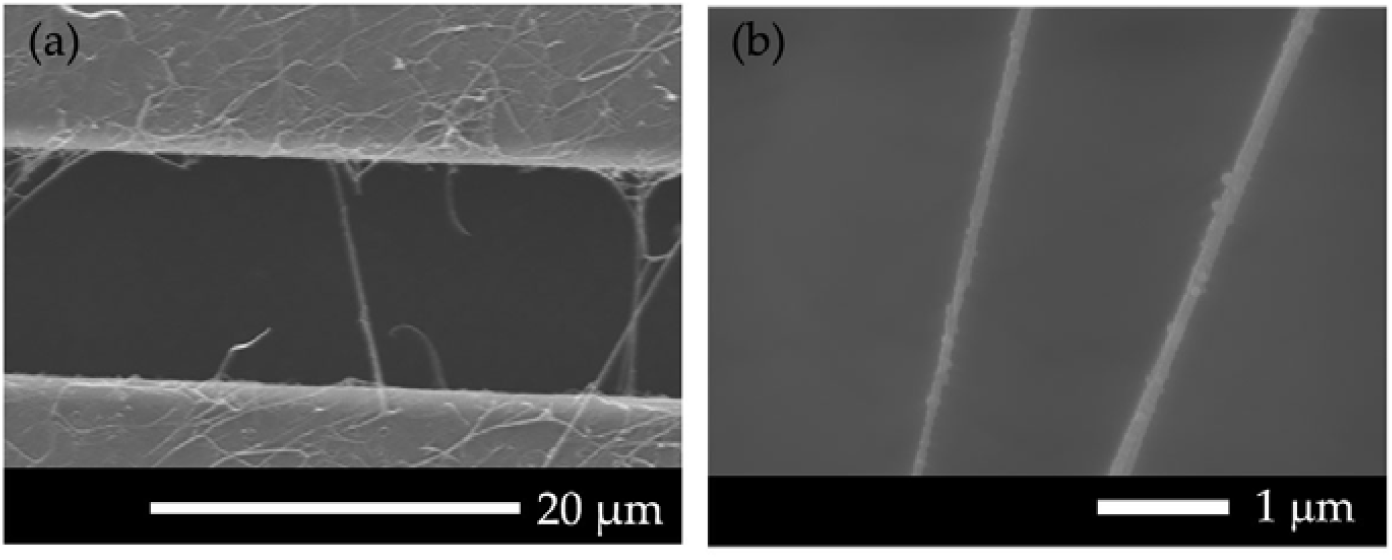
SEM images of fibrin fibers formed from 1 mg/ml fibrinogen at a) 900x and b) 23,000x magnification.

**Fig. 6.**
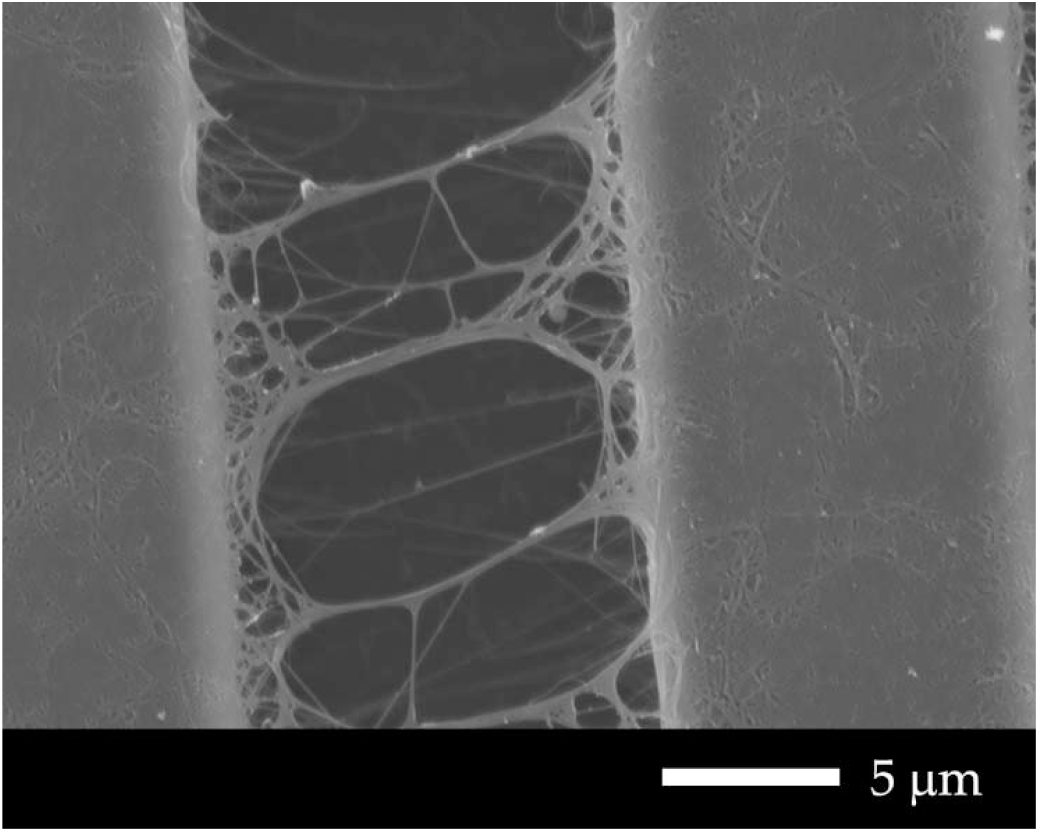
A SEM image at a magnification of 4,300x, of one population of fibrin fibers formed from 2 mg/ml fibrinogen. The image shows the striated pattern of the surfaces with fibers and film on top. Ridges can be seen on the right and left of the images and fibers span a 12 μm trough in between. The image shows the fibers originating from a root or film like structure at the ridge as opposed to continuing onto the ridge as seen in Fig. 5a.

#### 3.2.3 Fibrinogen concentration and initial modulus

Fibrin fibers are known to undergo modulus stiffening with increased strain. We measured the initial modulus, before stiffening, and the final modulus, after stiffening. The mechanical response of fibers formed from 1 mg/ml fibrinogen and 2 mg/ml fibrinogen were distinctly different from one another. On average, the initial modulus of fibers formed from 2 mg/ml fibrinogen was 10-times larger than the initial modulus of fibers formed from 1 mg/ml fibrinogen. The final modulus also followed the same trend with the final modulus of fibers formed from 2 mg/ml fibrinogen being significantly larger than those formed from 1 mg/ml fibrinogen. However, the final modulus of the 2 mg/ml fibrinogen fibers were only 3-times larger than the 1 mg/ml fibrinogen fibers. The initial and final modulus are reported in Table I.

**Table I.** The slope and modulus of fibers.

| Fibrinogen concentration (mg/ml) | Initial modulus (MPa) | Final Modulus (MPa) |
| --- | --- | --- |
| 1 | $3.9 \pm 1.2$ | $18.5 \pm 4.2$ |
| 2 | $39.0 \pm 12.0$ | $55.0 \pm 11.3$ |
| p-value | $p < 0.01$ | $p < 0.01$ |

Reported uncertainties are standard errors of the mean. n = 11 for the 1 mg/ml data and n = 15 the 2 mg/ml data

#### 3.2.4 Strain Hardening

As reported previously, fibrin fibers undergo strain hardening [21,42,47]. The stiffening of each fiber typically occurred over a small region of strain, enabling us to fit the stress strain curve before and after stiffening with two linear functions (Fig. 7). Interestingly not all fibers underwent modulus stiffening. All fibers formed from 1 mg/ml fibrinogen exhibited modulus hardening, while only 67 % of fibers formed from 2 mg/ml fibrinogen displayed modulus hardening. On average fibers formed from 1 mg/ml increased their modulus by a factor of 5.9 +/-1.1 (n = 11), whereas fibers formed from 2 mg/ml increased by a factor of 1.9 +/-0.2 (n = 15). Fig. 7 shows the strain hardening behavior and linear fit to determine the modulus of a typical fiber formed from 1 mg/ml and 2 mg/ml. Additionally, those formed with 1 mg/ml fibrinogen hardened at a significantly higher strain, 120 % +/-13 % compared to 60 % +/-6 % for fibers formed from 2 mg/ml (p < 0.01).

**Fig. 7.**
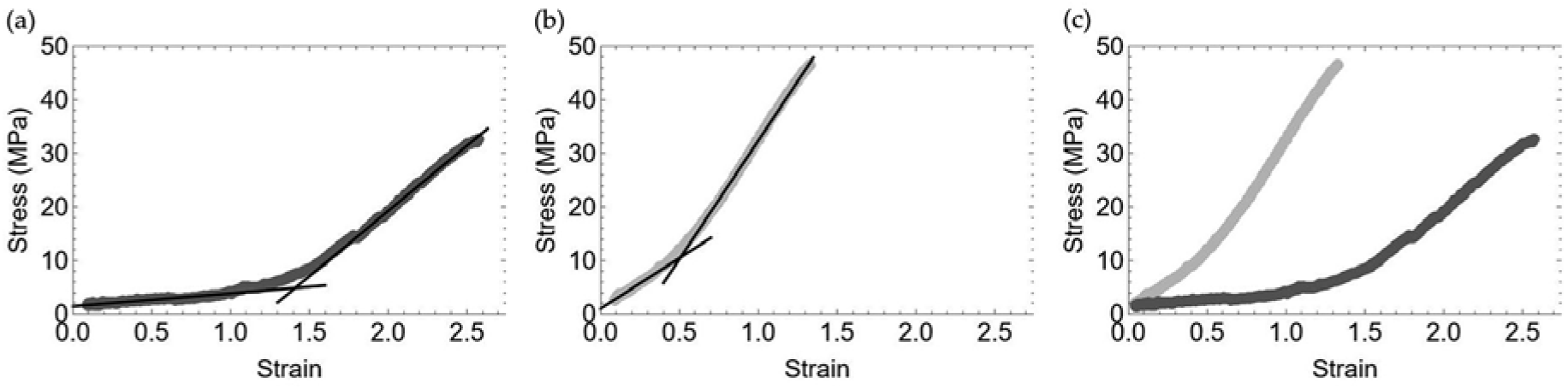
The stain hardening behavior of fibrin fibers formed from (a) 1 ml/ml fibrinogen and (b) 2 mg/ml fibrinogen. The initial and final slopes were fit with linear functions to obtain the moduli. (c) A representative graph comparing fibers formed from 1 mg/ml (dark gray) and 2 mg/ml fibrinogen (light gray). The fiber formed from 1 mg/ml is initially softer, undergoes strain hardening at a higher extensibility and is more extensible than its 1 mg/ml comparison. The fiber with a lower initial modulus does not reach the same maximum stress or force as its originally stiffer counterpart.

**Fig. 8.**
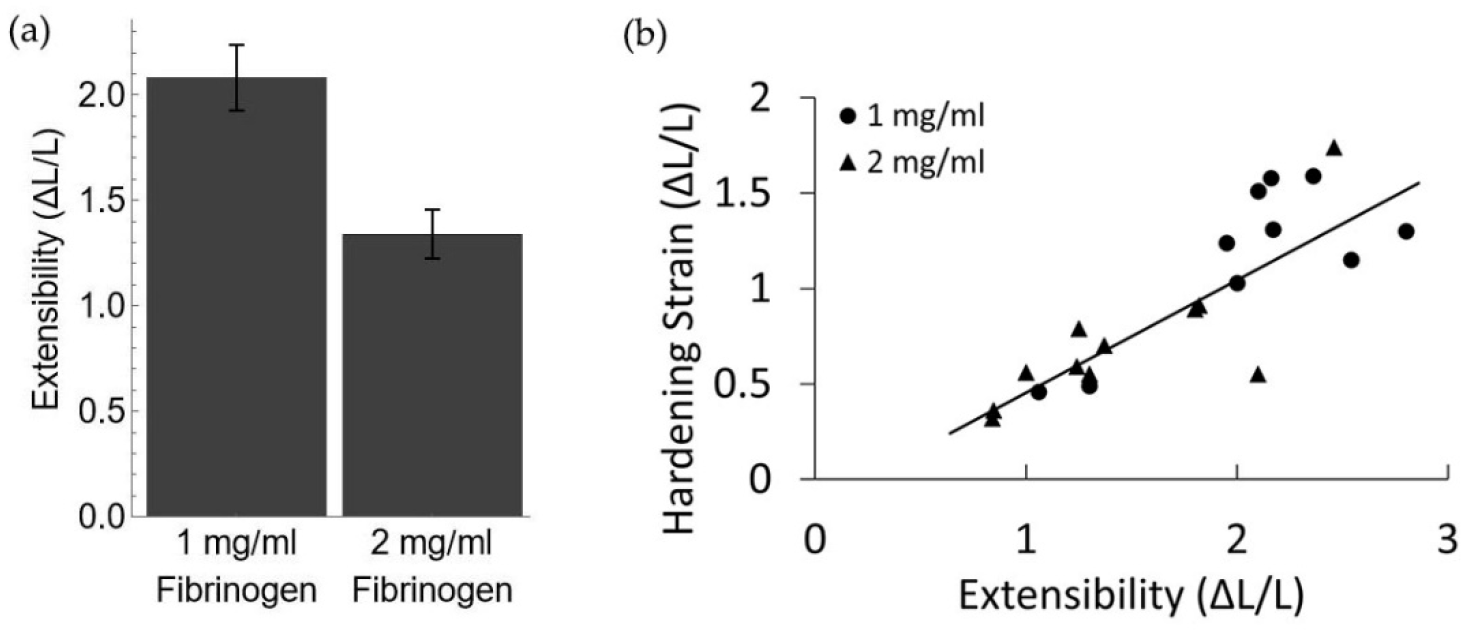
A) The average extensibility of fibrin fibers formed from 1 mg/ml and 2 mg/ml fibrinogen; error bars represent the standard deviation of the mean. B) A plot of the relationship between the strain at which the modulus increases, labeled hardening strain, and the maximum strain or extensibility of the fiber.

**Fig. 9.**
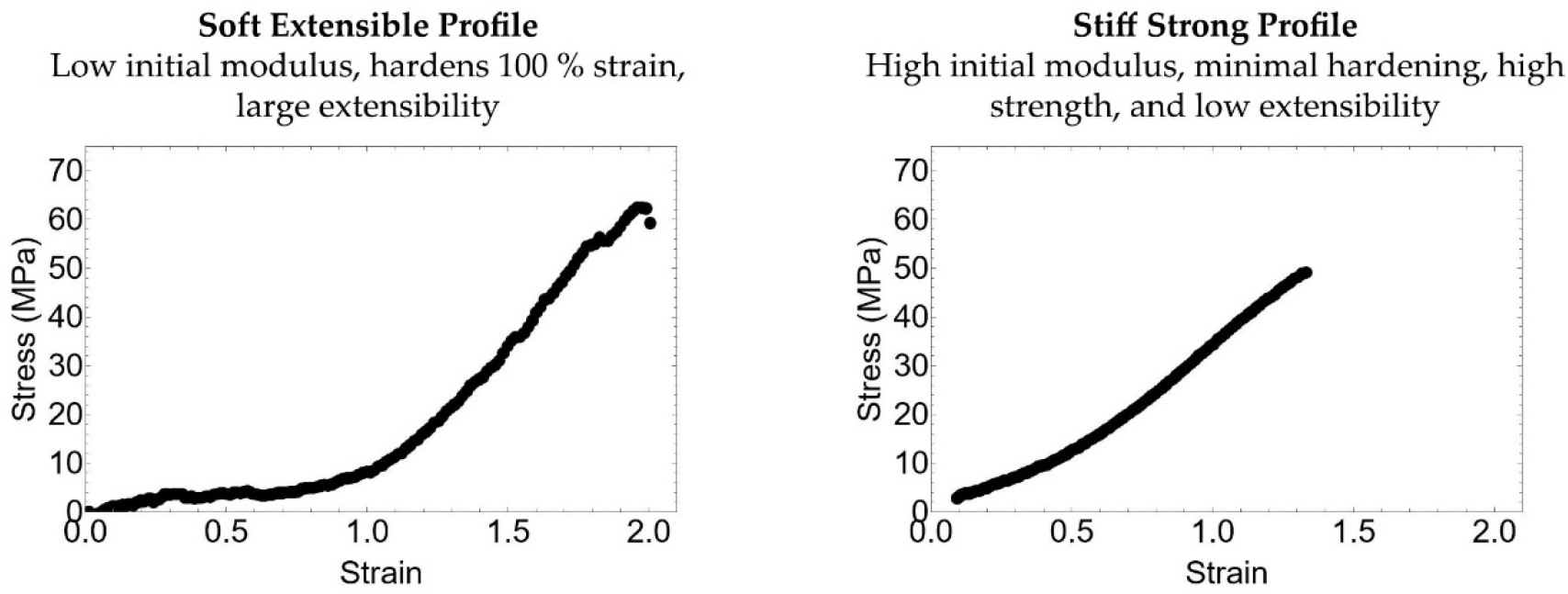
Representative stress-strain curves of fibers displaying the two fiber profiles.

#### 3.2.5 Extensibility

Fibers formed from 1 mg/ml had a significantly higher extensibility (p < 0.001). On average these fibers stretched to 208 % ± 16 % (n = 11) of their initial length while the average extensibility of fibers formed from 2 mg/ml fibrinogen was 134 % ± 12 % (n = 15). Analysis of fibers that undergo strain hardening showed a linear correlation between extensibility and the strain at which fibers harden. Fibers that waited to a higher strain to harden had a higher extensibility. Once the modulus hardened fibers formed from 1 mg/ml and 2 mg/ml fibrinogen extended an additional 86 % ± 9 % and 93 % ± 13 %, respectively, displaying no significantly different extension after the modulus increase.

#### 3.2.6 Maximum force

Fibers formed from 2 mg/ml fibrinogen ruptured at a higher applied force. This agrees with the results that 2 mg/ml fibers had a higher average force vs strain slope throughout manipulation. This increase in rupture force was significant from 190 nN +/-40 nN to 460 nN +/-120 nN (p < 0.05). Although the fibers with a lower initial slope displayed more significant hardening and extended to higher strains these properties did not raise the rupture force enough to match the rupture force of fibers with high initial slope (Fig. 7c).

## 4. Discussion

Individual fibrin fiber mechanical measurements show large variability within a fiber population. This is apparent when considering the reported standard deviations of the modulus of fibrin fibers which is often the same order of magnitude as the average. Additionally, there are differences in the reported average fibrin fiber extensibility. These variabilities both within a fiber population and between reported findings motivated our study of the variability of individual fibrin fiber mechanics.

Fibrinogen and thrombin in vivo naturally occur over a range of concentration. The approximate range of fibrinogen in plasma is from 2 mg/ml to 4 mg/ml and thrombin concentration depends on the time during the coagulation process and location within or adjacent to the site of coagulation [48]. Multiple research groups have shown that both fibrin concentration and thrombin concentration effect the rate of fiber formation and the resultant fiber diameter, length, branching, and clot density [49-51]. Trends in fiber extensibility are clear from aggregated published data. However, these trends do not depend on the ratio of thrombin to fibrinogen as much as the concentration of protein. This can be expected since the proportionality constant between protein concentration and structural clot changes differs for thrombin and fibrinogen [38]. We suspect the same can be said for the changes in fiber mechanics and chose to focus this investigation on the effects of fibrinogen concentration on fiber mechanics.

Because individual fiber extensibility is one of the easiest mechanical measurements to obtain, we used the extensibility results from a range of fibrinogen concentrations to guide our future data collection. The largest differences in fiber extensibility were found between the fibers formed from 1 mg/ml fibrinogen and 2 mg/ml fibrinogen. Therefore, we used these two concentrations for a more in-depth investigation into fibrin fiber mechanics.

In our purified fibrin system, we found significant differences in the mechanical properties of fibrin fibers formed from 2 mg/ml fibrinogen when compared to fibrin fibers formed form 1 mg/ml fibrinogen. Fibers formed from 2mg/ml had a higher modulus, were less extensible, and showed less strain hardening. While fibers formed from 1 mg/ml had a lower initial modulus, higher extensibility, and showed large strain hardening around 100 % strain. The modulus of 1 mg/ml fibers was initially 10 time lower than the 2 mg/ml fibrin fibers. However, after strain hardening the 1 mg/ml modulus was only 3 times lower than the 2 mg/ml modulus. Our data suggest the mechanism responsible for strain hardening is suppressed in fibers with a high initial modulus which was common among fibers formed from a higher fibrinogen concentration.

SEM images of the two samples may provide some insight into the cause for the mechanical differences. The SEM images showed fibers that continued onto the ridges as well as fibers with complex root like structures at the ridges. These complex structures resemble structures found by O’Brien et al. when first reporting fibrin film formation [9]. These root-like structures were seen predominately on the 2 mg/ml samples and could suggest a different formation process for many of the fibers on these samples. These stiffer non-hardening fibers may form from the rolling up and aggregation of fibrin film and therefore may lack the protofibril formation of traditional fibrin fibers [9]. This lack of protofibrils would eliminate extension via protofibril sliding which others speculate may cause hardening by an initial period of sliding protofibrils followed by limited motion due to αC interactions, therefore the lack of protofibrils could lead to a larger initial modulus and an absence of strain hardening. Interestingly, all fiber extended roughly 100 % strain beyond the point at which they strain hardened. This suggests the mechanism responsible for fiber extension is the same mechanism responsible for lower initial modulus.

Another, potential mechanism for the difference in fiber mechanics may be related to fibrin formation kinetic. The rate of protofibril formation and aggregation increases as fibrinogen concentration increases [38]. This faster polymerization may lead to differences in protein packing and protofibril interactions and therefore alter fiber mechanics.

The 1 mg/ml fibrinogen and 2 mg/ml fibrinogen fibers also showed differences in the force requires to rupture the fibers. The 2 mg/ml fibers on average ruptured at a higher force than the 1 mg/ml fibers. This is relevant to the physical role fibrin plays in stemming blood flow. It is reasonable to assume a lower rupture force would increase the chance of embolism.

The values presented above are the averages of all fibers formed at two different concentrations. When we consider each fiber independent of concentration the data revealed two distinct behavior profiles. The first profile is described by an initially soft fiber which undergoes modulus hardening around 100% strain or higher, ruptures at a high extensibility with a lower rupture force. The second profile is described by an initially stiff fiber that shows minimal modulus hardening occurring before 100 % strain, low extensibility, and a higher rupture force.

Fibers formed from 1 mg/ml fibrinogen were on average like the soft and extensible profile and had significantly lower moduli, higher extensibility, and ruptured at a lower stress than fibers formed from 2 mg/ml fibrinogen which on average were like the stiff and strong profile. However, there were fibers at both concentrations that more closely resembled the nondominant profile.

Our purified fibrin system is very different from a platelet poor plasma or whole blood system. It is unlikely that the behaviors of each concentration, i.e. 1 mg/ml and 2 mg/ml, relate directly to those same concentrations in the more complex system. However, it is possible and likely that the variation in fibrin mechanics measured in the purified system would also be present in the more complicated system.

## 5. Conclusion

This work shows the natural variations in fibrin fiber mechanics. It describes two distinct mechanical behavior profiles seen in multiple individual fibers. Clots comprised of fibers with the soft extensible profile would respond to external stress in a very different manner than clots formed from the stiff strong profile. The behavior of fibers in a clot would affect clot contraction, extensibility, and risk of embolism. This work elucidates these differences and generates many questions about the source of these variation in fiber mechanics. For example, does fibrin film formation play a role? Are both behavior profiles found in vivo? What is the roll of protofibril sliding in fiber mechanics and how can we alter protofibril sliding? Continued study of the mechanics of fibrin fibers and clots will help to deepen our understanding of mechanisms responsible for these differences and hemostasis and thrombosis.

## Data Availability

Analyzed data are contained within the article. The data that support the findings of this study are available on request from the corresponding author, CCH, upon reasonable request. Script for controlling the MFP-3D-BIO AFM is openly available at https://facultystaff.richmond.edu/~chelms/publications.html.

## Acknowledgements

The authors would like to acknowledge the University of Richmond Faculty Research Fellowships.

